# TandemMapper and TandemQUAST: mapping long reads and assessing/improving assembly quality in extra-long tandem repeats

**DOI:** 10.1101/2019.12.23.887158

**Authors:** Alla Mikheenko, Andrey V. Bzikadze, Alexey Gurevich, Karen H. Miga, Pavel A. Pevzner

**Affiliations:** Center for Algorithmic Biotechnology, Institute of Translational Biomedicine, Saint Petersburg State University, Saint Petersburg, Russia; Graduate Program in Bioinformatics and Systems Biology, University of California, San Diego, CA, USA; UC Santa Cruz Genomics Institute, University of California, Santa Cruz, CA, USA; Department of Computer Science and Engineering, University of California, San Diego, CA, USA

## Abstract

Extra-long tandem repeats (ETRs) are widespread in eukaryotic genomes and play an important role in fundamental cellular processes, such as chromosome segregation. Although emerging long-read technologies have enabled ETR assemblies, the accuracy of such assemblies is difficult to evaluate since there is no standard tool for their quality assessment. Moreover, since the mapping of long error-prone reads to ETR remains an open problem, it is not clear how to polish draft ETR assemblies. To address these problems, we developed the tandemMapper tool for mapping reads to ETRs and the tandemQUAST tool for polishing ETR assemblies and their quality assessment. We demonstrate that tandemQUAST not only reveals errors in and evaluates ETR assemblies, but also improves them. To illustrate how tandemMapper and tandemQUAST work, we apply them to recently generated assemblies of human centromeres.

## Introduction

Tandem repeats are formed by multiple consecutive nearly identical sequences that are often generated by unequal crossover (Smith, 1976). The early DNA sequencing projects revealed that tandem repeats are abundant in eukaryotic genomes (Yunis and Yasmineh, 1971; Bacolla et al., 2008). Recent studies of tandem repeats revealed their role in various cellular processes and demonstrated that mutations in tandem repeats may lead to genetic disorders (McFarland et al., 2015; Giunta and Funabiki, 2017; Song et al., 2018; Black et al., 2018).

We distinguish between extensively studied short tandem repeats (Willems et al. 2014; Gymrek et al., 2016; Saini et al., 2018) and *extra-long tandem repeats* (ETRs) that range in length from tens of thousands to millions of nucleotides. Since ETRs are difficult to assemble, the vast majority of them remain unassembled even in the human genome, let alone other species. Centromeres and pericentromeres contain some of the longest ETRs, that account for ∼3% of the human genome and span megabase-long regions (Miga, 2019). They represent the “dark matter” of the human genome that evaded all attempts to sequence it so far and are the biggest gaps in the reference human genome (Hayden et al., 2013; Miga et al., 2019).

Emergence of long-read technologies, such as Pacific Biosciences (PacBio) and Oxford Nanopore Technologies (ONT), have greatly altered the landscape of whole-genome sequencing. The development of long-read assemblers (Chin et al., 2016; Koren et al., 2017; Kolmogorov et al., 2019; Ruan and Li, 2019) and hybrid assemblers that combine long and short reads (Antipov et al., 2016; Zimin et al., 2017) significantly increased the contiguity of assembled genomes compared to short-read assemblies. In addition, long reads contributed to successful semi-manual approaches for reconstructing human centromeres (Jain et al., 2018a; Miga et al., 2019). The Flye assembler successfully resolves *bridged tandem repeats* that are spanned by long reads and even some *unbridged tandem repeats* that are not spanned by long reads (Kolmogorov et al., 2019). The centroFlye assembler (Bzikadze and Pevzner, 2019) was designed to automatically assemble unbridged ETRs, such as centromeres.

Various alternative strategies for ETR assembly and absence of the ground truth for benchmarking these assemblies raise the problem of their quality evaluation. Similar problems have been addressed by the short-read quality assessment tools for genome assemblies such as GAGE (Salzberg et al., 2011) and QUAST (Gurevich et al., 2013; Mikheenko et al., 2018) as well as specialized quality assessment tools metaQUAST (Mikheenko et al., 2016) and rnaQUAST (Bushmanova et al., 2016). However, these tools are based on known references and thus are not applicable to analyzing ETRs since their analysis requires *reference-free* approaches to evaluating assembly quality. At the same time, existing reference-free tools are based on analyzing paired-end read alignments or gene content (Hunt et al. 2013; Clark et al. 2013; Ghodsi et al. 2013; Simão et al. 2015) and are not applicable to ETRs either.

Existing assembly quality assessment tools rely on aligners (Li and Durbin, 2009; Langmead et al., 2009; Li, 2016; Li, 2018) that accurately map reads to assemblies. However, our benchmarking revealed that these tools often fail in ETRs, for example, minimap2 (Li, 2018) results in incorrect alignments of some reads to ETRs, especially in regions with assembly errors. We thus developed the tandemMapper tool that efficiently maps long error-prone reads to ETRs. TandemMapper not only enabled tandemQUAST development but also led to improvement in ETR assemblies due to more accurate read mapping and subsequent polishing.

The initial attempt to evaluate the quality of ETR assemblies was centromere-specific (Bzikadze and Pevzner, 2019) and has not resulted in a general quality assessment tool for ETR assemblies. Species- and chromosome-specific nature of centromeres prevents application of the same approach to other ETRs. However, the common principles of centromere organization can be utilized for developing a universal assembly evaluation tool for ETRs.

Centromeres of primates are comprised of retrotransposon repeats and AT-rich *alpha satellites*, a DNA repeat based on a 171 bp monomer (Manuelidis and Wu, 1978). In humans and many primates, consecutive monomers are arranged tandemly into *higher-order repeat* (*HOR*) *units* (Willard and Waye, 1987a). The number of monomers and their order in the HOR are chromosome-specific. For example, the chromosome X HOR, referred to as DXZ1, consists of twelve monomers (Willard and Waye, 1987b). These twelve monomers evolved from an ancestral pentameric satellite repeat ABCDE and can be represented as C_1_D_1_E_1_ A_1_B_1_C_2_D_2_E_2_A_2_B_2_C_3_D_3_. For consistency with Bzikadze and Pevzner, 2019, we took the liberty to refer to the chromosome X HOR as ABCDEFGHIKL.

Here we present tandemMapper, a tool for mapping reads to ETRs, and tandemQUAST, a tool for evaluating and improving ETR assemblies. We used tandemMapper and subsequent polishing to modify assemblies of the human centromere X generated by both centroFlye (Bzikadze and Pevzner, 2019) and curated semi-manual approach (Miga et al., 2019). We further illustrated tandemQUAST work by analyzing quality of resulting assemblies and demonstrating that they improve on original assemblies. These improvements suggest that tandemQUAST will become a popular tool for evaluating quality and polishing of many assemblies since nearly all genomes have ETRs.

TandemMapper and tandemQUAST are open-source software that are freely available as command-line utilities on GitHub at https://github.com/ablab/tandemQUAST.

## Methods

As an input, tandemQUAST requires one or several ETR assemblies and the set of long reads (PacBio continuous long reads (CLR) or ONT) that contributed to these assemblies. Additionally, error-prone long reads can be complemented by accurate reads such as PacBio high-fidelity (HiFi) reads (we do not consider accurate but short Illumina reads since they proved to be very difficult to unambiguously map to a centromere). TandemQUAST consists of the read mapping module that identifies positions of read alignments to the assembly, polishing module for improving the quality of an assembly based on the identified read alignments, and the quality assessment module. TandemQUAST uses *general metrics* for evaluating ETRs of any kind and *centromeric metrics* designed specially to account for HOR structure of centromeric ETR.

### Simulated assembly

To evaluate tandemMapper and tandemQUAST results, we simulated an ETR of length ∼1.03Mb which is a concatenation of 500 randomly mutated copies of the consensus HOR sequence on chromosome X (DXZ1) that diverge from the consensus sequence by 1% (substitutions only). Then, we simulated 1400 reads from this ETR using NanoSim (Yang et al., 2017) trained on the real ONT dataset enriched for ultra-long reads (longer than 50 kb) and generated by Telomere-to-Telomere (T2T) consortium (Miga et al., 2019). We refer to the centroFlye assembly of these reads as *simulated*. We further introduced various artificial errors (described below) into the simulated assembly and ran tandemQUAST.

### TandemMapper module

The key part of many long-read assemblers is a read mapping procedure that operates with short sequences of length *k* or simply *k-mers*. Most long-read mapping algorithms are based on *minimizers* (Li, 2016; Jain, 2018b; Li, 2018), *k-*mers that are carefully chosen and used as stepping stones for read mapping. However, mapping a long read to an ETR is a non-trivial problem since minimizers are expected to be reduced in numbers and irregularly arranged due to local expansions of identical tandem repeats. Bzikadze and Pevzner, 2019 used *unique k-mers* (that appear just once in the assembly) to improve read mapping to ETRs. However, the T2TX7 assembly of chromosome X (referred to as cenX) has only 16,163 unique 21-mers across the 3.1 Mb cenX array, with the largest distance between unique 21-mers equal to 42 kb (Miga et al., 2019).

The density of unique *k*-mers may significantly vary along an assembly, leading to incorrect mappings and drops in coverage by mapped reads in some regions (Figure 1). Therefore, tandemMapper uses *locally unique k-mers* that are more abundant than unique *k*-mers. It partitions the assembly into *t* segments (the default value *t*=5) and defines a *locally unique k-mer as* a *k*-mer that is unique in a given segment. The segment size may vary depending on the assembly length, read lengths, and distribution of unique *k*-mers along the assembly. Figure 1 illustrates that density of locally unique *k*-mers is significantly larger than density of unique *k*-mers, thus providing more “signposts” for read mapping.

**Figure 1.**
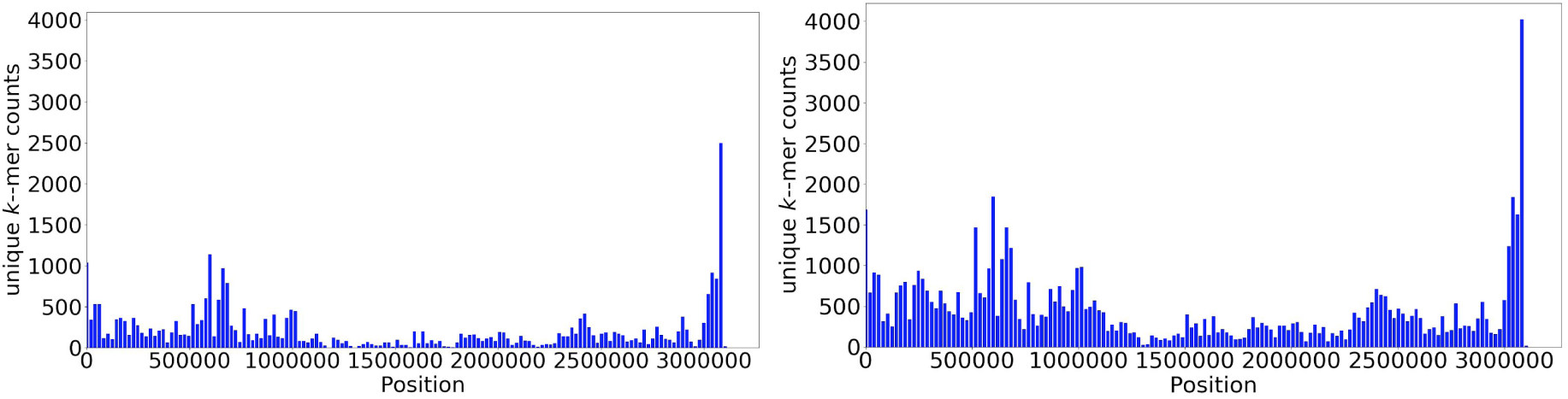
Distribution of unique (left) and locally unique (right) *k*-mers along the cenX assembly of the CHM13 cell line constructed by centroFlye. Each bar shows the number of unique (locally unique) *k*-mers in a bin of length 20 kb. The total number of unique (locally unique) *k*-mers is 33,545 (41,296) with *k*=19 and *t*=5 (each segment is approximately 600 kb long).

Since ETR assemblies can be error-prone, some locally unique *k*-mers may represent assembly errors rather than low-frequency *k*-mers in the genome. To filter out such locally unique *k*-mers we analyze their frequencies in the read set. We assume that a *k*-mer from an assembly was erroneously attributed to locally unique if it has an unusually low frequency (lower than *MinFrequency*) or an unusually high frequency (higher than *MaxFrequency*) in reads. The *MinFrequency* (*MaxFrequency)* threshold is defined as a value such that at least 1% (95%) of all locally unique *k*-mers have the same or lower frequency in reads.

We select locally unique *k*-mers that occur in reads at least *MinFrequency* and at most *MaxFrequency* times, and refer to them as *solid k-mers.* Filtering *k*-mers by frequency in reads reduces the number of spurious *k*-mers erroneously defined as locally unique. For example, applying this filtration to the centroFlye cenX assembly v0.8.3 (Bzikadze and Pevzner, 2019) reduced the number of locally unique *k*-mers from 41,296 to 37,728. Comparison with PacBio HiFi reads generated from the same cell line (Vollger et al., 2019) revealed that 1,723 of 3,586 filtered out *k*-mers are absent in the HiFi read set or, on the contrary, have a very high frequency (higher than a frequency of 95% of *k*-mers in the read set).

The *k*-mer selection procedure can be affected by the fact that ETRs may harbor various transposable elements (*TEs*) such as LINE repeats, Alu repeats, etc. Single copies of TEs within ETRs are likely to contain many locally unique *k*-mers that may affect the mapping accuracy and complicate further analysis. To minimize their influence, we mask TEs using RepeatMasker (Smit and Green, http://repeatmasker.org) before selecting locally unique *k*-mers.

The tandemMapper algorithm is inspired by the minimap2 (Li, 2018) and Flye mappers (Lin et al., 2016, Kolmogorov et al., 2019). Given two solid *k*-mers *a* and *b*, shared between a read *R* and an assembly *A*, we define *d*_*R*_*(a,b)* and *d*_*A*_*(a,b)* as distances between *a* and *b* in *R* and *A*, respectively. We further define *distance(a,b)=min*{*d*_*R*_*(a,b),d*_*A*_*(a,b)*}, *diff(a,b)=|d*_*R*_*(a,b)-d*_*A*_*(a,b)*|, and *penalty(a,b)=diff(a,b)/distance(a,b).* We call *k*-mers *a* and *b compatible* if *distance(a,b)* < *maxDistance* (default value *maxDistance =* 60 kb) and *diff*(*a,b*) < *C * distance*(*a,b*), where *C* is a constant (the default value is 0.15).

Given a read, we define a directed weighted *compatibility graph* with a vertex set equal to a set of all solid *k*-mers shared between *R* and *A*. We connect vertices *a* and *b* by an edge if (i) *a* precedes *b* in *R* and (ii) *a* and *b* are compatible. We further define the weight of an edge between *a* and *b* as *premium*-*penalty(a,b)*, where *premium* is a constant selected to optimize the number of correctly mapped reads (default value *penalty*=0.1). A *chain* between *R* and *A* is defined as the longest path in the compatibility graph.

A chain for a given read represents a potential mapping of this read to the assembly. TandemMapper finds a chain for each read using dynamic programming, filters out short chains (shorter than 3 kb in length or containing less than 20 *k*-mers by default), and constructs the corresponding nucleotide alignments within the derived chain boundaries for each remaining chain.

We benchmarked tandemMapper and minimap2 by aligning simulated reads to the *simulated* assembly and comparing their known exact positions in the assembly to the inferred positions (Table 1). To analyze how these metrics capture breakpoints, we generated *simulated*_*del*_ assembly by introducing an artificial deletion of length 10 kb in the *simulated* assembly at position 400 kb.

**Table 1.** Benchmarking of tandemMapper and minimap2 on the simulated dataset. A read is considered correctly mapped if its starting position is within 100 bp from the read start position calculated for the longest read alignment (an alignment is elongated to both ends of a read). Only alignments longer than 3 kb were considered.

| | tandemMapper<br>(unique $k$-mers) | tandemMapper<br>(locally unique $k$-mers) | minimap2 |
| --- | --- | --- | --- |
| # mapped reads | 1228 | 1242 | 1239 |
| # incorrectly mapped reads | 4 | 2 | 34 |
| # reads spanning the deletion breakpoint | 0 | 0 | 58 |

TandemMapper split all alignments spanning the breakpoint of this deletion, while minimap2 erroneously extended alignments through this breakpoint due to highly repetitive sequence of the ETR. Using locally unique *k*-mers instead of unique *k*-mers increased the number of correctly mapped reads even in an easy case of the simulated assembly with uniform density of distribution of unique *k*-mers.

### Polishing module

Due to high error rate in reads, most long-read assemblers have to include a polishing step to improve base-calling accuracy of the assembly (Chin et al., 2013; Loman et al., 2015; Lin et al., 2016). However, our benchmarking revealed that standard polishing tools may even decrease the assembly quality in tandem repeats due to incorrect and ambiguous read alignments against the assembly. On the other hand, Miga et al., 2019 demonstrated that the *marker-assisted read mapping* (based on unique *k*-mers) significantly improves accuracy of ETR assemblies. TandemQUAST uses read alignments generated by tandemMapper as an input for a modified Flye polishing module (Lin et al., 2016, Kolmogorov et al., 2019). The Results section demonstrates that this polishing procedure fixes erroneous deletions and base-calling errors.

### Quality assessment module

To evaluate the assembly quality and reveal possible errors, we developed two *general* metrics (indel-based and *k*-mer-based) and a *centromeric* metrics (monomer-based) that we describe below. General metrics are applicable to any ETRs and centromeric metrics are applicable to centromeric ETRs only.

### Indel-based metrics

ETR assemblies are prone to large-scale deletions and duplications that lead to *misassembly breakpoints*. QUAST (Gurevich *et al.*, 2013) defines a misassembly breakpoint based on differences between an assembly and a reference genome. In contrast, since the reference is not available, tandemQUAST detects breakpoints based on abnormalities in read coverage. Below we describe the *coverage* metric and the *breakpoint* metric and use them to reveal putative breakpoints. To analyze how these metrics capture breakpoints, we used the *simulated*_*del*_ assembly (Figure 2).

**Figure 2.**
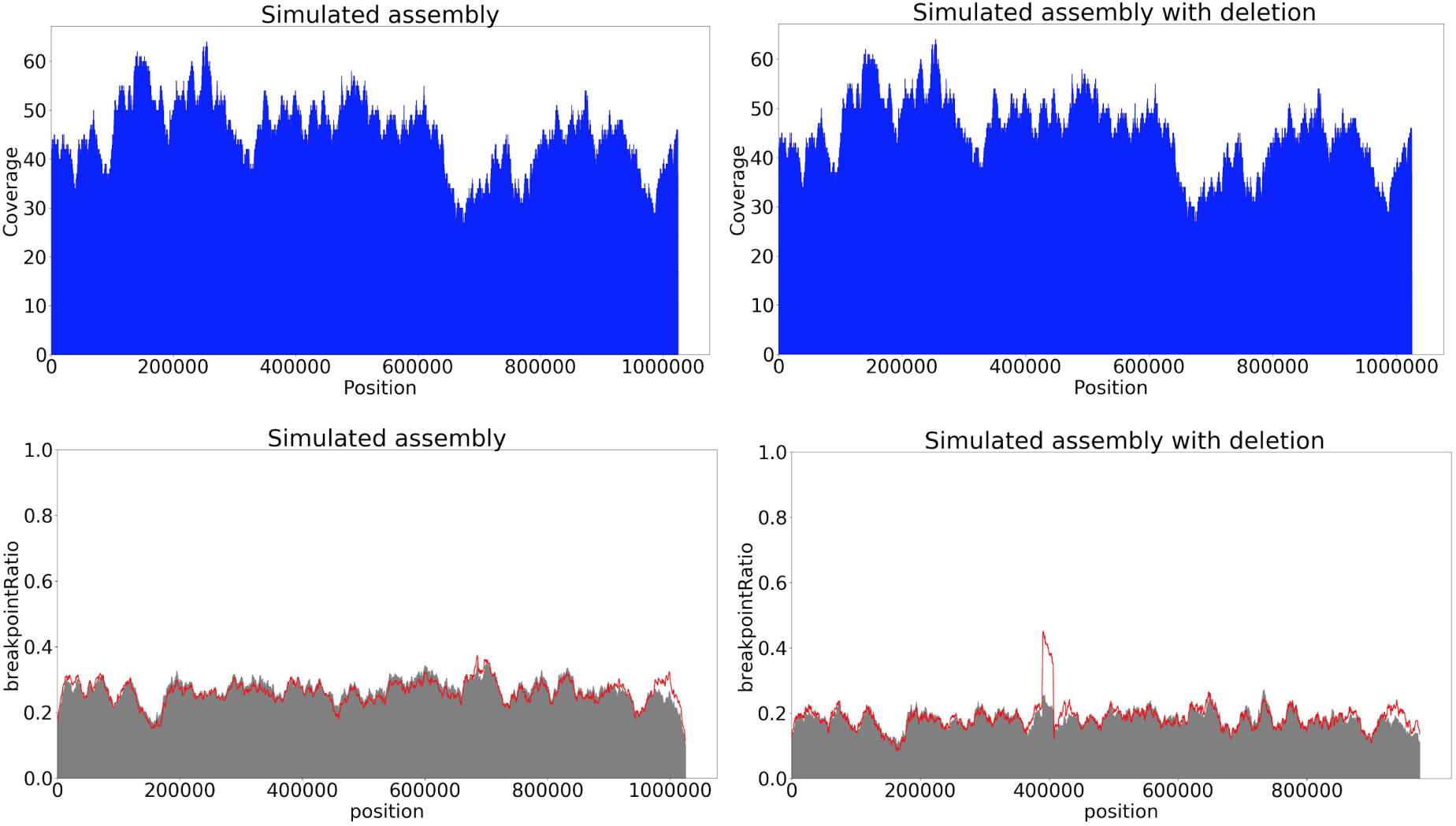
Coverage (top) and breakpoint (bottom) metrics for *simulated* (left) and *simulated*_*del*_ (right) assemblies. The coverage plot does not show a significant drop at the point of the deletion but the breakpoint plot reveals a peak at the position of the deletion (400 kb). The red plot is based on the *breakpointRatio(Kmer)* values, the gray plot is based on the *breakpointRatio*^*+*^*(Kmer)* values.

#### Coverage metric

Assembly errors may affect the coverage near the assembly breakpoints. TandemQUAST uses read mappings truncated with respect to their longest chains to construct the coverage plot and reveal regions with abnormal coverage that may point to assembly errors (Figure 2).

#### Breakpoint metric

In case an assembly contains a breakpoint caused by a long indel, longest chains for the majority of reads spanning this indel breakpoint cannot be extended through this indel due to a substantial discrepancy in distances between solid *k*-mers in reads and the assembly. Thus, if longest chains for many reads start or end in a certain region, this region may contain an assembly breakpoint.

However, stochastic differences in coverage and various biases also may result in drops or peaks in read coverage. Our goal is to distinguish these cases and reveal assembly breakpoints.

A chain for a read *R* defines its partitioning into *prefix*(*R*), *middle*(*R*), *and suffix*(*R*), where *middle*(*R)* starts at the first *k*-mer in the chain and ends in the last *k*-mer in the chain. This chain defines a *chain-segment* in the assembly between the first and the last *k*-mer in the chain that is aligned to *middle*(*R*). We also define an *elongated chain-segment* as a chain-segment extended by |*prefix*(*R*)*|* and *|suffix*(*R*)*|* nucleotides in the beginning and the end, respectively.

Given a solid *k*-mer *Kmer*, we define *breaks*(*Kmer)* as the number of chains starting or ending in this *k*-mer (over all reads). We also define *number(Kmer)* (*number*^*+*^*(Kmer)*) as the number of chain-segments (elongated chain-segments) containing this *k*-mer. Finally, we define *breakpointRatio*(*Kmer*) as *breaks*(*Kmer)/number(Kmer)* and *breakpointRatio*^*+*^(*Kmer*) as *breaks*(*Kmer)/number*^*+*^(*Kmer*).

While drops in values of *breakpointRatio* usually correspond to poorly covered regions, peaks in values may reveal breakpoints in the assembly. We expect that regions where *breakpointRatio(Kmer)* has significantly higher values than *breakpointRatio*^*+*^*(Kmer)* contain assembly breakpoints, because the longest chains for many reads were not extended through this region (Figure 2).

### *k*-mer-based metrics

To benchmark metrics evaluating the base-calling accuracy of an assembly, we introduced 10,000 (∼1% of the sequence length) random single-nucleotide substitutions in the *simulated* assembly (we refer to this assembly as *simulated*_*mut*_).

In contrast to the tandemMapper tool, the *k*-mer-based metrics need a reliable set of *k*-mers that appear just once in the assembly. We thus filter out solid *k*-mers that occur more than once in the assembly or more than once in a single read and refer to the rest as *unique solid k-mers*.

After constructing read alignments, tandemQUAST finds where a unique solid *k*-mer in a read aligns to the assembly and calculates coordinates of all found alignments across all reads containing this *k*-mer. Afterwards, it clusters these coordinates (for a given unique solid *k*-mer) if they are located within *MaxClumpDistance* from each other (default value *MaxClumpDistance* = 1 kb). After single linkage clustering, we define a cluster as a *clump* if it contains more than *MinClumpSize* elements (default value *MinClumpSize* = 2). Ideally, all occurrences of a unique solid *k*-mer should form a single clump. We divide all *k*-mers having at least *MinClumpSize* occurrences in reads into three groups: a single clump, multiple clumps, and spurious *k*-mers that do not form clumps (Figure 3). TandemQUAST reports the percentage of each group and their distribution in the assembly (Figure 4).

**Figure 3.**
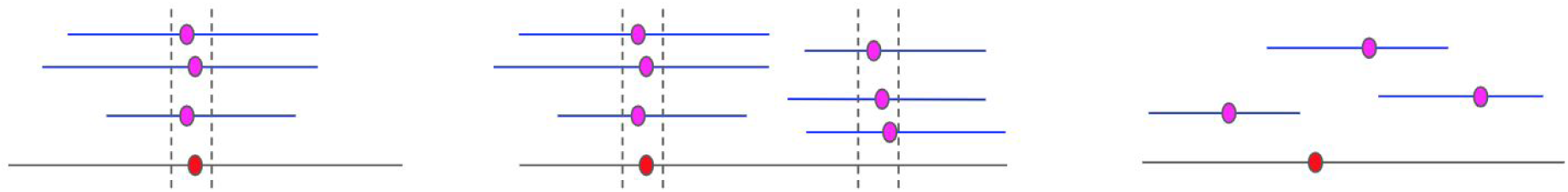
Coordinates of unique solid *k*-mers in the assembly and reads. Purple and red dots represent *k*-mer position in reads (shown as blue lines) and in the assembly (shown as a gray line), respectively. Clumps are flanked by vertical lines. (Left) *k*-mers forming a single clump, (Middle) *k*-mers forming multiple clumps in different parts of the assembly, (Right) *k*-mers that do not form clumps (spurious *k-*mers).

**Figure 4.**
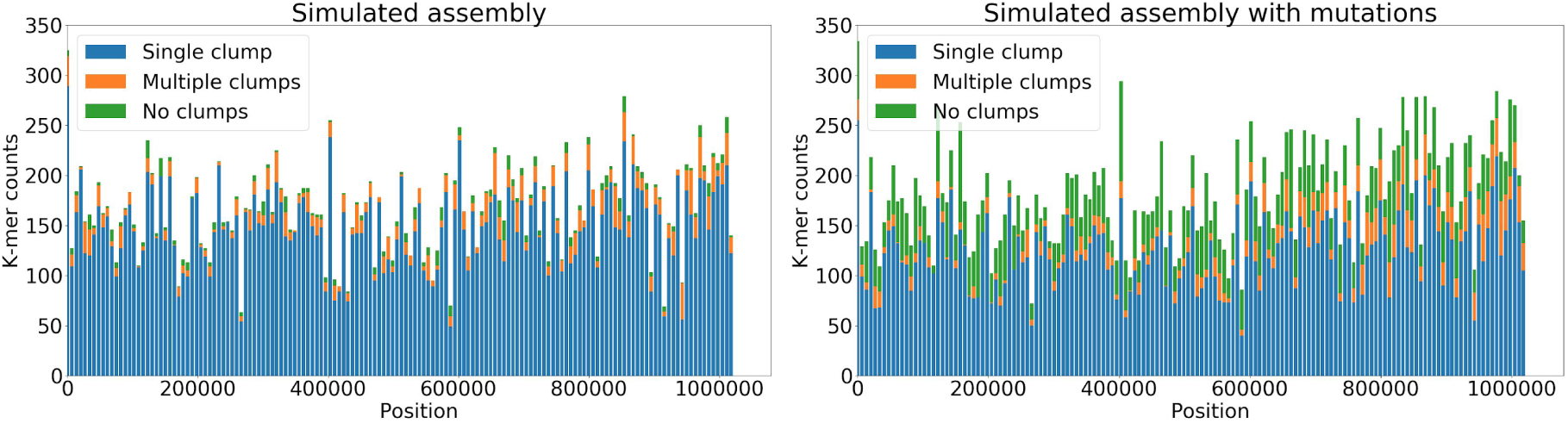
Distribution of different types of unique solid *k*-mers in the simulated (left) and *simulated*_*mut*_ (right) assemblies. Each bar shows the number of different types of *k*-mers in a bin of length 5 kb. The total number of unique *k*-mers in the assembly that do not occur in reads increased from 3,269 in the *simulated* assembly to 13,792 in the *simulated*_*mut*_ assembly. The percent of unique solid *k*-mers forming a single clump decreased from 91% in the *simulated* assembly to 74% in the *simulated*_*mut*_ assembly, mostly due to increased number of spurious *k-*mers.

In the case when a complementary set of accurate PacBio HiFi reads is provided, tandemQUAST compares *k*-mer frequencies in the assembly and accurate reads. If the assembly contains *k*-mers that do not occur in HiFi reads or frequent *k*-mers from reads have a low frequency or are even absent in the assembly, it is likely that the assembly requires additional polishing. TandemQUAST reports absolute and relative abundance of such *k*-mers and generates a plot showing their distribution (Figure 11 in the Results section). Multiple clumps or spurious *k*-mers appearing along the entire assembly may point to poor base-calling quality of this assembly. Multiple clumps or spurious *k*-mers appearing in certain regions of an assembly reflect either a poor base-calling quality in these regions or collapsed duplications with subsequent “consensus” polishing with reads from both copies.

### Centromeric metrics

The additional set of metrics takes into account centromere organization into monomers and units. Currently, tandemQUAST focuses on analysis of a particular type of centromeres that are formed by HORs. When a set of specific monomer sequences is known, tandemQUAST can analyse the assembly using the *monomer-based* metric described below and the *unit-based* statistic described in Appendix “Unit-based statistic”. In order to illustrate monomer-based metric and unit-based statistic, we generated the *simulated*_*del_monomer*_ assembly by introducing a deletion of 3 consecutive monomers in the *simulated* assembly at position 226 kb.

Centromere assemblies may include difficult-to-detect indels of multiple monomers. In case monomer sequences are known, tandemQUAST attempts to detect discrepancies between reads and the assembly at the monomer level. The assembled centromere and all reads are aligned to the provided monomer sequences and are subsequently translated into the monomer alphabet using the StringDecomposer tool (Dvorkina et al., 2019), resulting in a *monocentromere* and *monoreads*. Using nucleotide read alignments, for each monomer *ReadMonomer* in each monoread tandemQUAST calculates *StartPos(ReadMonomer)*, the starting nucleotide position of *ReadMonomer* in the monocentromere. In case *ReadMonomer* is aligned against a deletion in the monocentromere, *StartPos(ReadMonomer)* is recursively defined as *StartPos(NextReadMonomer)* where *NextReadMonomer* is the following monomer in the monoread. For each monomer *CenMonomer* in the monocentromere we define *StartPos(CenMonomer)* as the starting nucleotide position of this monomer in the centromere. We define *ReadMonomers(CenMonomer)* as a multiset of such *ReadMonomers* that *|StartPos(ReadMonomer) - StartPos(CenMonomer)|* < *MaxStartPosDist* (the default value *MaxStartPosDist* = 50 bp). Finally, we define *MonomerRatio(CenMonomer)* as the frequency of *CenMonomer* in *ReadMonomers(CenMonomer)*. If *MonomerRatio(CenMonomer)* is below *MinMonomerRatio* (default value *MinMonomerRatio* = 0.8), the assembly is likely to have an error (Figure 5). However, in the case of heterozygous sites this ratio is close to 0.5 as roughly half of the reads support (do not support) the monomer.

**Figure 5.**
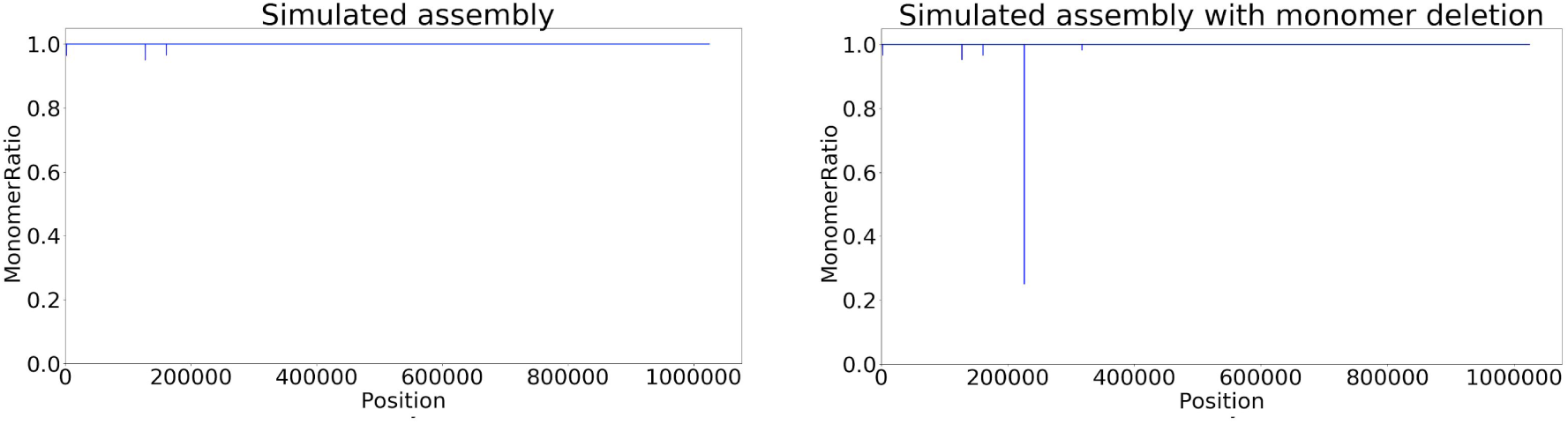
*MonomerRatio* for *simulated* and *simulated*_*del_monomer*_ assemblies. Even though *MonomerRatio* is defined for *CenMonomers*, we prefer to show nucleotide coordinates over the centromere (X-axis) for the sake of consistency with other metrics. The sharp drop in *MonomerRatio* in the *simulated*_*del_monomer*_ assembly corresponds to the position of the monomer deletion.

Although individual monomers may significantly vary in sequence, their length is fairly conserved within species that have alpha-satellites (Haaf and Willard, 1998; Hall et al., 2003). Thus, the monomer length distribution across the centromere assembly in such species may point to flaws in the assembly. Figure 6 demonstrates that most monomers have conserved length across the assembly. However, the first monomer A and the last monomer L show surprising variability in length, suggesting that the accuracy of the simulated assembly deteriorates at the ends of HOR units due to imperfect polishing. This imperfect polishing is caused by limitations of the existing read mapping tools in ETRs, forcing centroFlye to perform separate polishing for each HOR. Since the polishing procedure (Lin et al., 2016) is known to have limitations in the very beginning/end of each segment subjected to polishing, the beginning of the first (A) and the end of the last (L) monomers are poorly polished.

**Figure 6.**
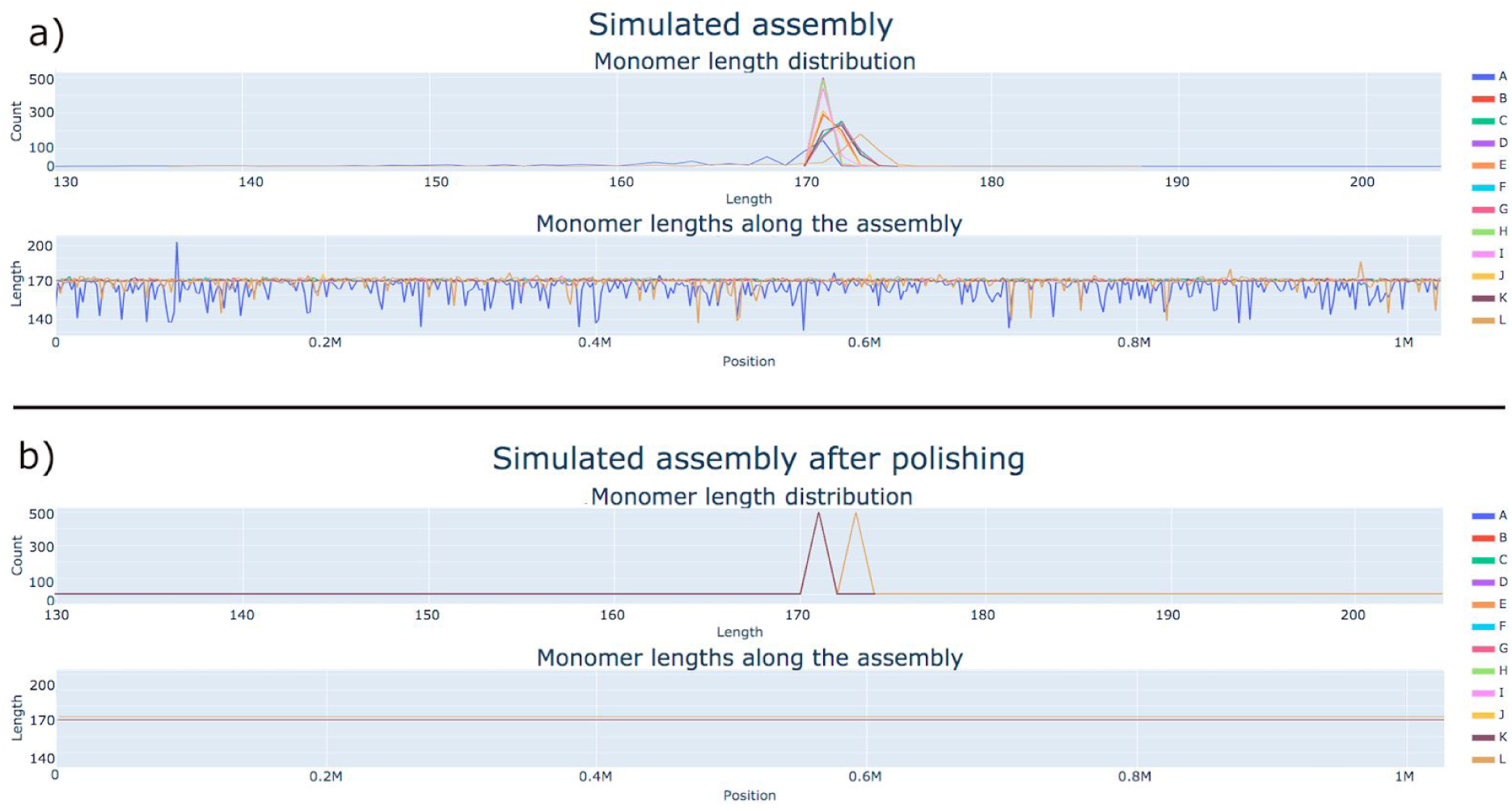
Monomer length distribution for the *simulated* (a) and s*imulated*_*polish*_ (b) assemblies. Monomer sequences forming a consensus DXZ1* sequence derived in Bzikadze and Pevzner, 2019 were used for analysis. In the *simulated* assembly, the length of A-monomers varies from 131 to 203 bp (mean 165 bp) and the length of L-monomers varies from 137 to 187 bp (mean 171 bp). In the s*imulated*_*polish*_ assembly, the length of all A-monomers (L-monomers) is equal to 171 (173) bp. Since all monomers, except for L, have lengths 171 bp after polishing, they all are represented by the color corresponding to the K-monomer.

Since tandemMapper accurately maps reads, it eliminates the need to polish each HOR separately and thus improves polishing of the first and the last monomers. Just a single round of polishing with tandemQUAST resulted in the s*imulated*_*polish*_ assembly with increased assembly length (by ∼4 kb) and complete sequences of the first and last monomers (Figure 6).

### Comparison of various ETR assemblies

TandemQUAST performs pairwise comparison for each pair of analyzed assemblies using the *bi-mapping* plot and the *discordance* test.

A bi-mapping plot (Figure 7) provides an overview of read alignments from the perspective of both assemblies. Each read aligned to both assemblies represents a dot with its starting mapping positions in two assemblies as the *x*- and *y*-coordinates. Positions of read alignments for two assemblies can be compared to reveal structural discrepancies between them.

**Figure 7.**
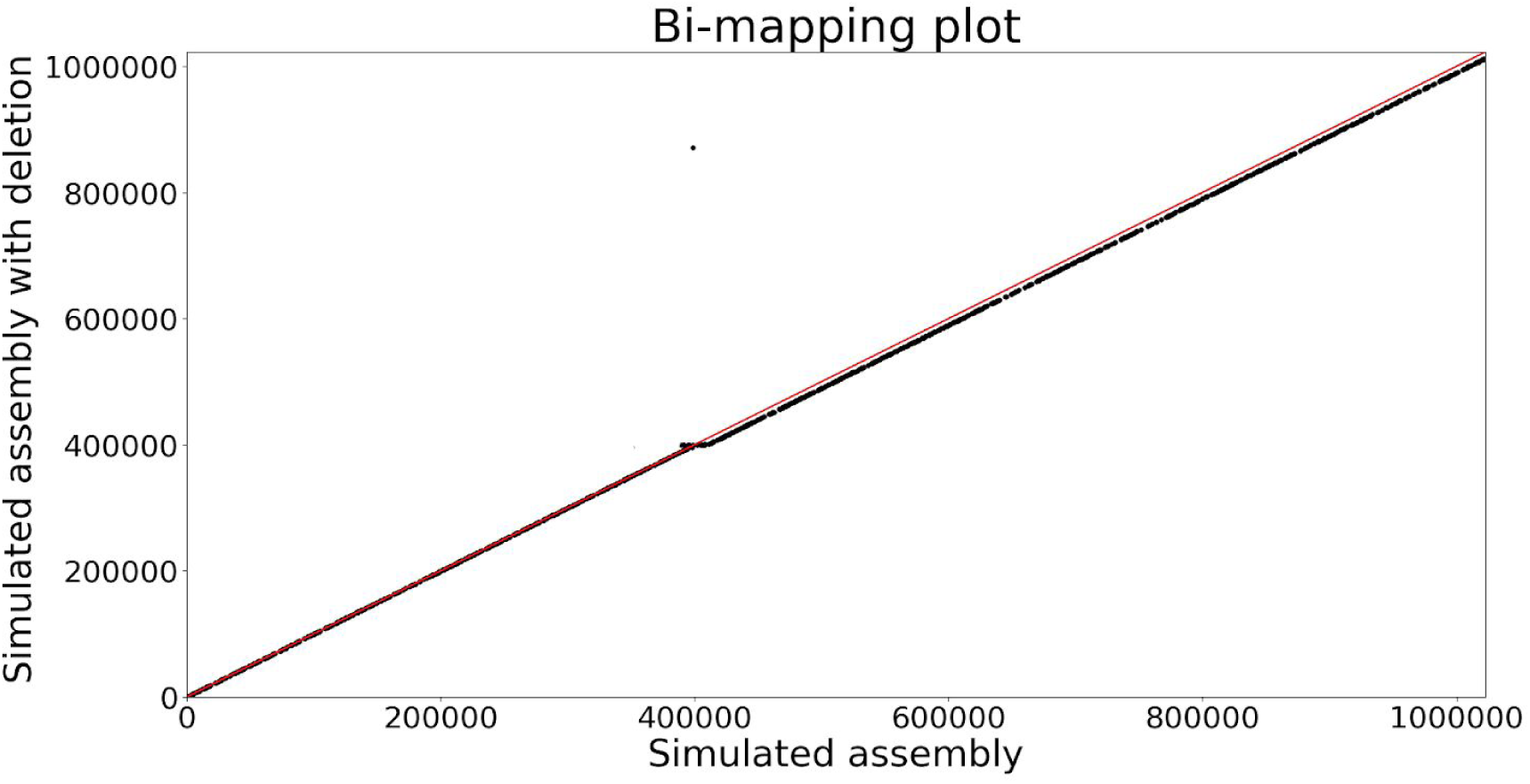
The bi-mapping plot illustrates the discrepancy between *simulated* and *simulated*_*del*_ assemblies at the deletion breakpoint.

Bzikadze and Pevzner, 2019 introduced the discordance test for comparing two assemblies. A *k*-mer is shared between an assembly and a read aligned to this assembly if it occurs in both the assembly and the read approximately at the same position in their alignment. Given a set of *k*-mers *Anchors*, we define *sharedAnchors*(*Read, Assembly*) as the number of *k*-mers from *Anchors* that are shared between *Read* and *Assembly*. The larger *sharedAnchors*(*Read, Assembly*) is, the better the assembly “explains” the read with respect to a given set of *k*-mers. Given a read set *Reads*, we define *sharedAnchors(Reads, Assembly)* as the sum of *sharedAnchors(Read, Assembly)* over all reads in *Reads*.

To compare two assemblies, we define *Anchors* as the set of shared unique *k*-mers between them (the default value *k*=19) and compute the discordance between these assemblies as *discordance(Assembly’, Assembly’’)* = *sharedAnchors(Reads, Assembly’)* - *sharedAnchors(Reads, Assembly’’)*. We classify a read *Read* as *discordant* with respect to assemblies *Assembly’* and *Assembly’’* and a set of *k*-mers *Anchors* if there is a large difference (by at least *k*) between *sharedAnchors(Read, Assembly’)* and *sharedAnchors(Read, Assembly’’)*, thus showing preference for one of the assemblies. We say that a discordant read *votes* for *Assembly’ (Assembly’’)* if this difference is positive (negative).

Figure 8 shows a cluster of discordant reads voting for *simulated* over *simulated*_*del*_ assembly at the deletion breakpoint and no reads voting for *simulated*_*del*_ assembly.

**Figure 8.**
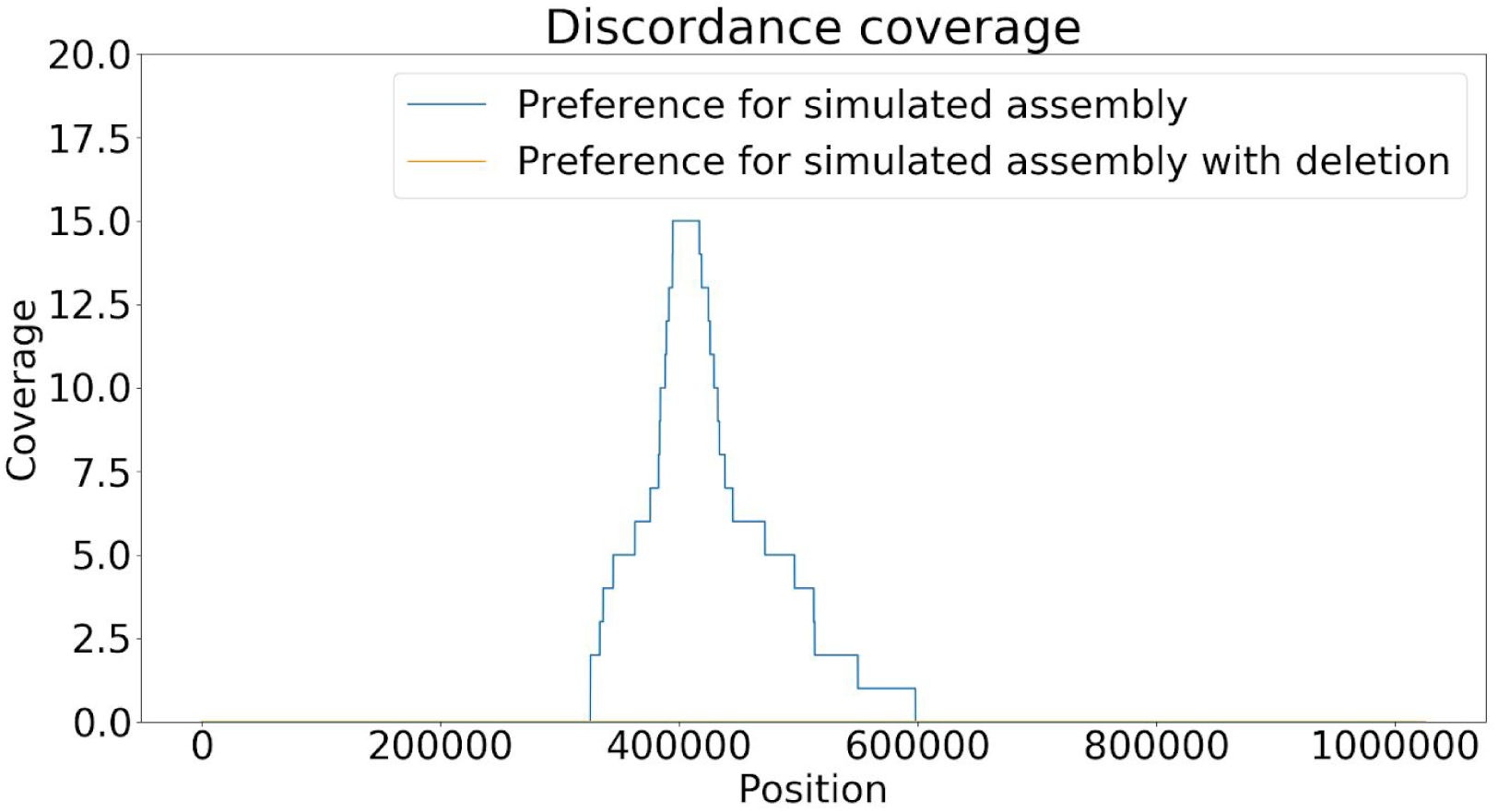
Coverage of *simulated* and *simulated*_*del*_ assemblies by discordant reads.

## Results

### Analysis of cenX assemblies

We analyzed the following centromere X (cenX) assemblies: the Telomere-to-Telomere consortium assembly v0.4 (T2T4), v0.7 (T2T7) (Miga et al., 2019), and centroFlye v0.8.3 assembly (centroFlye) (Bzikadze and Pevzner, 2019). Note, that the T2T4 assembly is an interim version that was not polished with the marker-assisted methods described in Miga et al., 2019. We added it to the comparison to show how tandemQUAST analyzes unpolished assemblies. The T2T7 version was first semi-manually assembled and further improved based on centroFlye assembly as described in Miga et al., 2019. The T2T7 assembly was further polished using a novel marker-assisted read mapping strategy using both nanopore and PacBio CLR reads. In contrast, the centroFlye assembly utilized only information derived from ONT reads at the polishing step.

We also applied our polishing method to the T2T4 and centroFlye assemblies (resulting in T2T4_polish_ and centroFlye_polish_ assemblies) to demonstrate how tandemQUAST improves assemblies.

### Indel-based metrics

Figure 9 illustrates that T2T4, T2T4_polish_, and centroFlye assemblies have a coverage drop in the center of the centromere at ∼1300-1600 kb that has a low concentration of unique *k*-mers (Figure 10).

**Figure 9.**
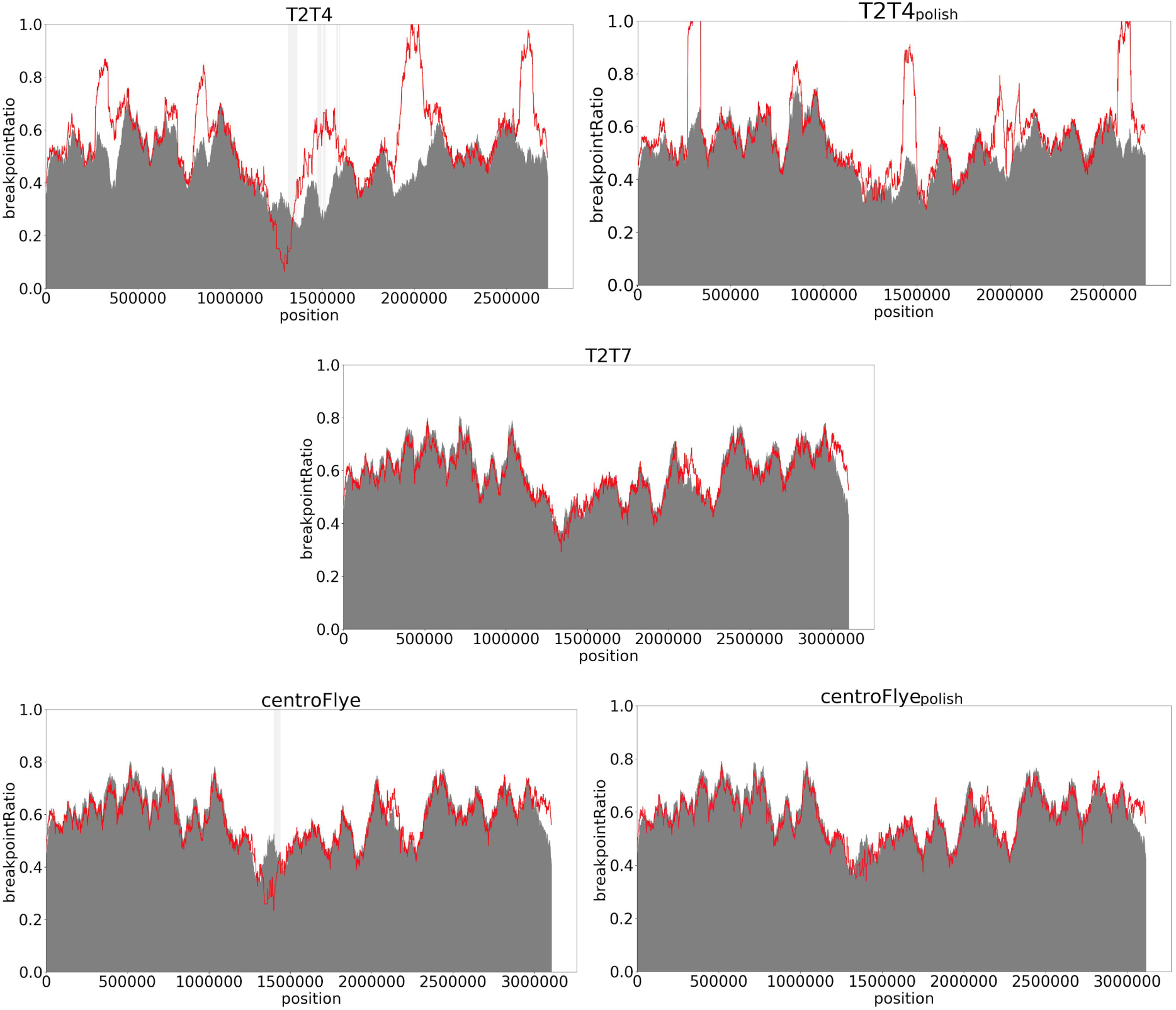
Breakpoint metric for the T2T4, T2T4_polish_, T2T7, centroFlye, and centroFlye_polish_ assemblies. The red plot and the gray plot are based on the *breakpointRatio(Kmer) and breakpointRatio*^*+*^*(Kmer)* values correspondingly. The vertical light gray bands represent regions with low coverage (<10x). Discrepancies in these regions should be considered as not necessarily related to flaws in an assembly.

**Figure 10.**
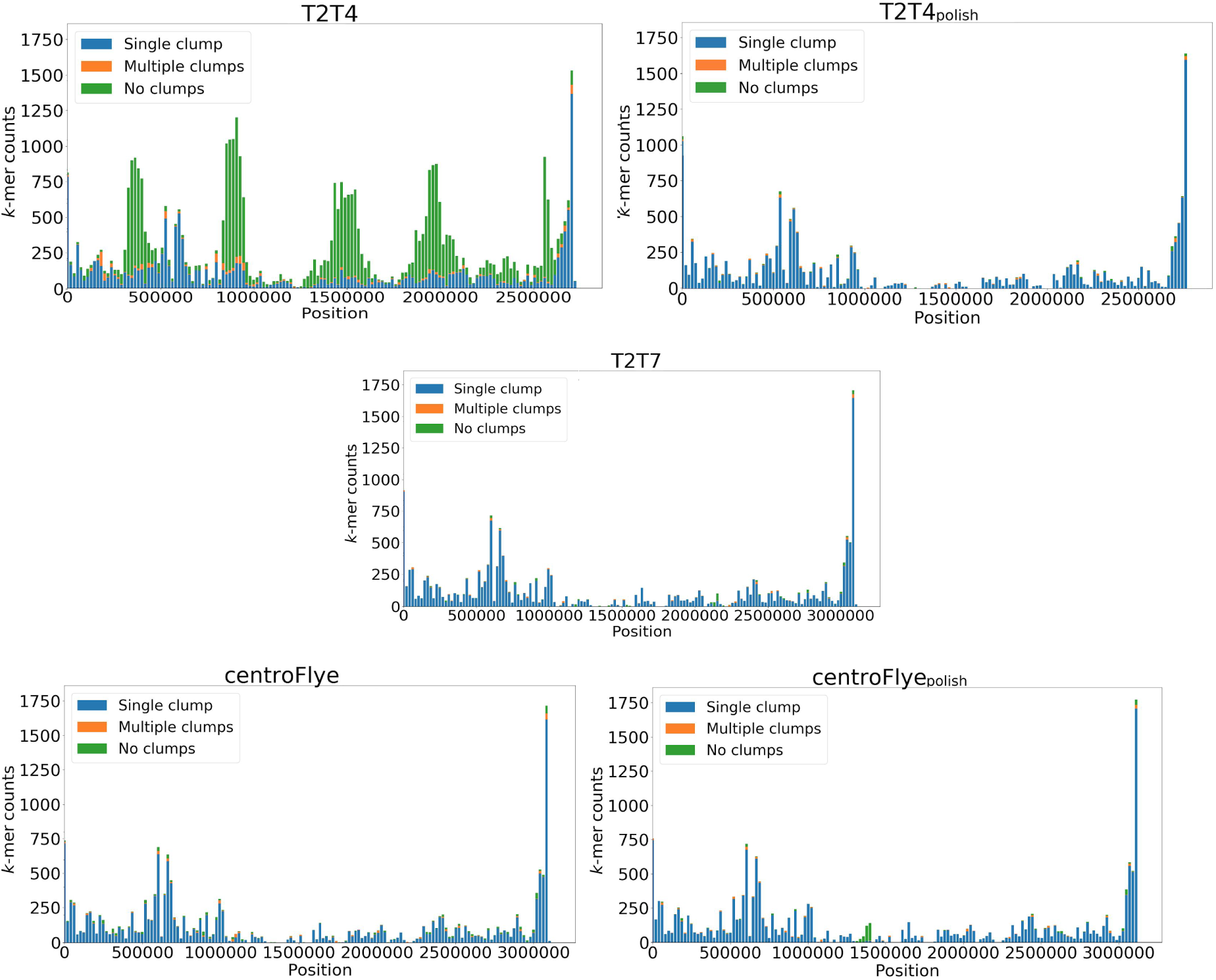
Distribution of different types of unique solid *k*-mers along the T2T4, T2T4_polish_, T2T7, centroFlye, and centroFlye_polish_ assemblies. Each bar shows the number of different types of *k*-mers in a bin of length 20 kb. The green peaks in the T2T4 assembly show that most unique solid *k*-mers in the assembly are spurious due to limited polishing.

**Figure 11.**
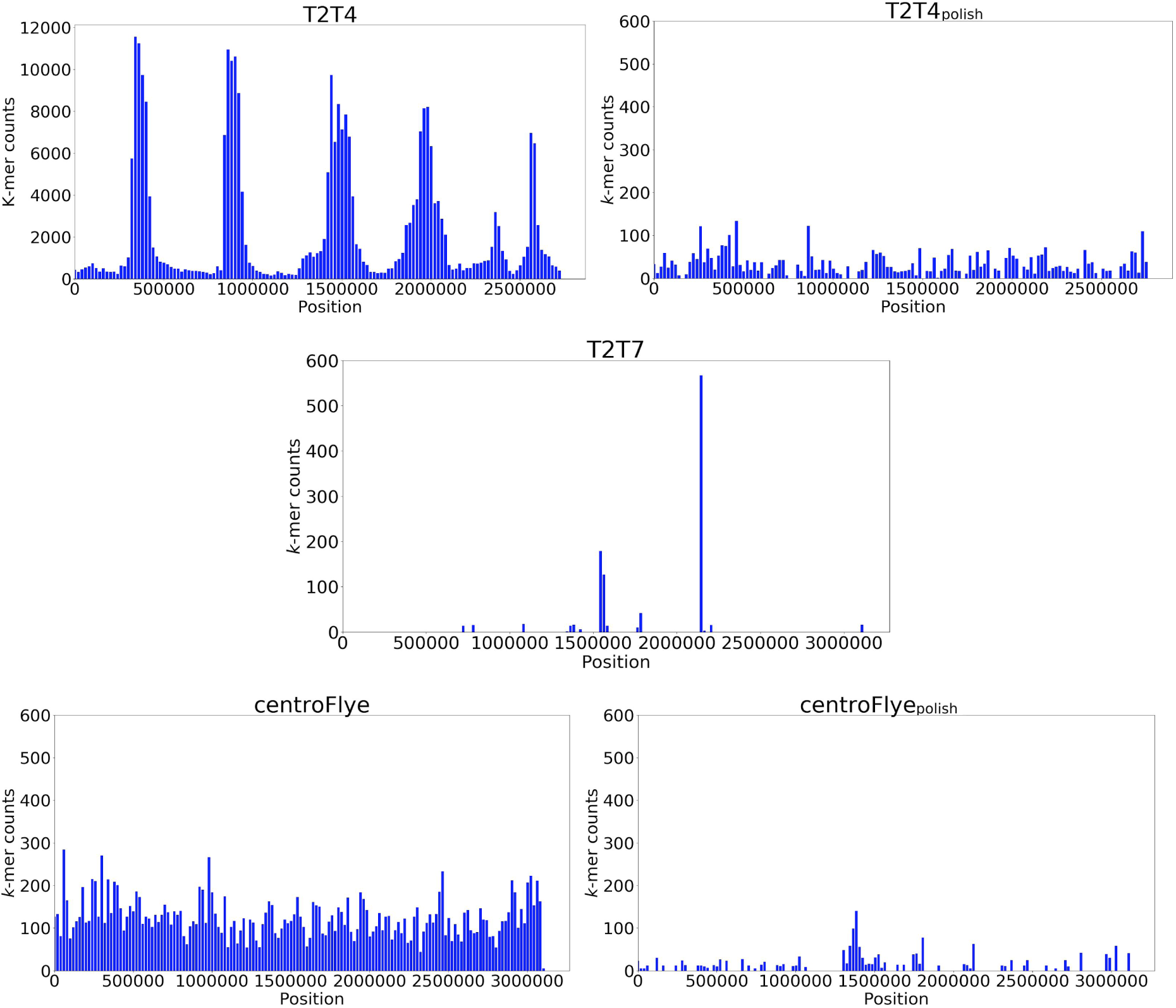
Distribution of *k*-mers absent in PacBio HiFi read set but present in the T2T4, T2T4_polish_, T2T7, centroFlye, and centroFlye_polish_ assemblies. Each bar shows the number of *k*-mers in a bin of length 20 kb that are present in an assembly but missing in HiFi reads. The numbers of *k*-mers that do not occur in the HiFi read set are 223,579 (T2T4), 5,038 (T2T4_polish_), 842 (T2T7), 7,867 (centroFlye), and 1,284 (centroFlye_polish_).

Low base-calling accuracy of the assembly can prevent chain extension. As a result, the longest chains for many reads may end in a poorly polished region, causing an increase in *breakpointRatio* values. Thus, to verify breakpoints found in the T2T4 assembly, we compared them to the T2T4_polish_ assembly. Both assemblies have peaks in *breakpointRatio* values at ∼270 kb, ∼800 kb, ∼1500 kb, ∼2000 kb, and ∼2500 kb that correlate with their bi-mapping plot (Figure 13). A small peak at ∼800kb reveals a deletion (∼3.5kb) in T2T4 and T2T4_polish_. The breakpoint metric for centroFlye and T2T7 assemblies are generally consistent between *breakpointRatio(Kmer)* and *breakpointRatio*^*+*^*(Kmer)* values, suggesting that these assemblies do not have large indels and rearrangements.

### *k*-mer-based metric

Figure 10 and Table 2 show the distribution of different types of unique solid *k*-mers across the assemblies. The T2T4 assembly has a very high number of spurious *k*-mers as expected for an unpolished assembly, while T2T4_polish_ demonstrates significant improvement in base-calling accuracy across the assembly. The high percentage (92-96%) of *k*-mers forming a single clump in the T2T7 and centroFlye assemblies suggest a high base-level quality in these assemblies.

**Table 2.** Distribution of different types of unique solid *k*-mers in the T2T4, T2T4_polish_, T2T7, centroFlye, and centroFlye_polish_ assemblies. Most *k*-mers forming multiple clumps form clumps of size 2. If we set *MinClumpSize* = 3, only 31 *k*-mers form multiple clumps and only 16 of them are in non-overlapping positions. Note that T2T4, T2T7, centroFlye and centroFlye_polish_ assemblies do not utilize information derived from accurate HiFi PacBio reads.

|  | T2T4 | T2T4<sub>polish</sub> | T2T7 | centroFlye | centroFlye<sub>polish</sub> |
| --- | --- | --- | --- | --- | --- |
| single clump | 14848<br>(36%) | 16004<br>(95%) | 16276<br>(96%) | 15956<br>(92%) | 16732<br>(95%) |
| multiple clumps | 1566<br>(4%) | 496<br>(3%) | 351<br>(2%) | 513<br>(3%) | 423<br>(2%) |
| no clumps | 24814<br>(60%) | 284<br>(2%) | 363<br>(2%) | 929<br>(5%) | 628<br>(3%) |

In addition, we compared *k*-mer frequencies in assemblies and in accurate PacBio HiFi reads generated from the same cell line CHM13 (Vollger et al., 2019). The number of *k*-mers that do not occur in the HiFi read set was the highest in the unpolished T2T4 assembly (223,579) and the lowest (842) in the T2T7 assembly.

### Monomer metrics

Figure 12 presents the monomer length distribution across various assemblies. The T2T7 and centroFlye assemblies have a few unusually short (145-146 bp) A-monomers at ∼1000 kb. We checked these monomers further and confirmed that they are supported by reads. Besides that, the T2T7 assembly has very conserved monomer lengths except for a few monomers at ∼2150 kb. In the centroFlye assembly, L-monomers significantly vary in length as in the simulated assembly (Figure 6), suggesting that centroFlye assembly requires additional polishing of HOR unit ends. The centroFlye_polish_ assembly has significantly more uniform monomer lengths as compared to the centroFlye assembly.

**Figure 12.**
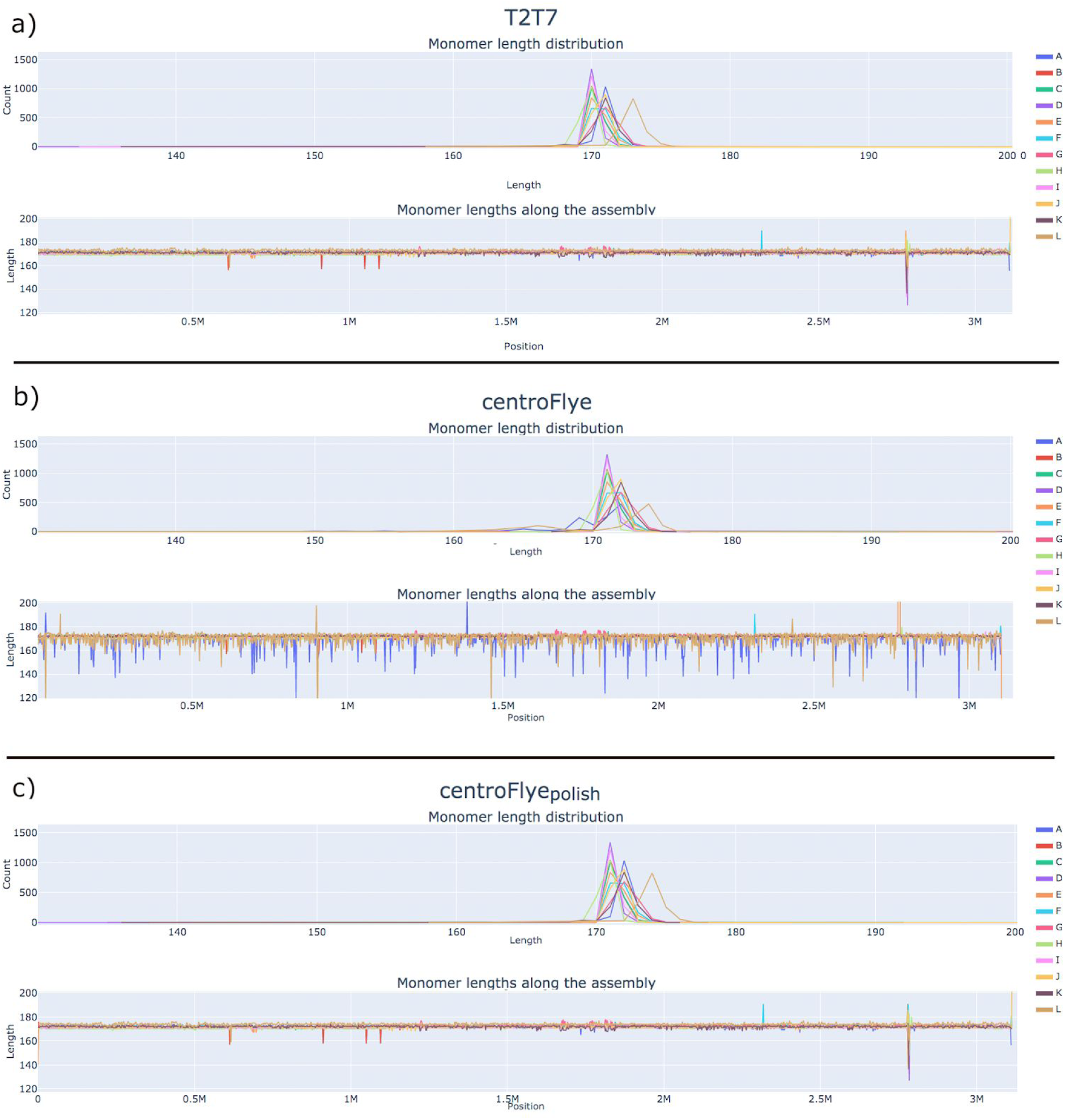
Monomer length distribution along the assembly in the T2T7, centroFlye, and centroFlye_polish_ assemblies.

**Figure 13.**
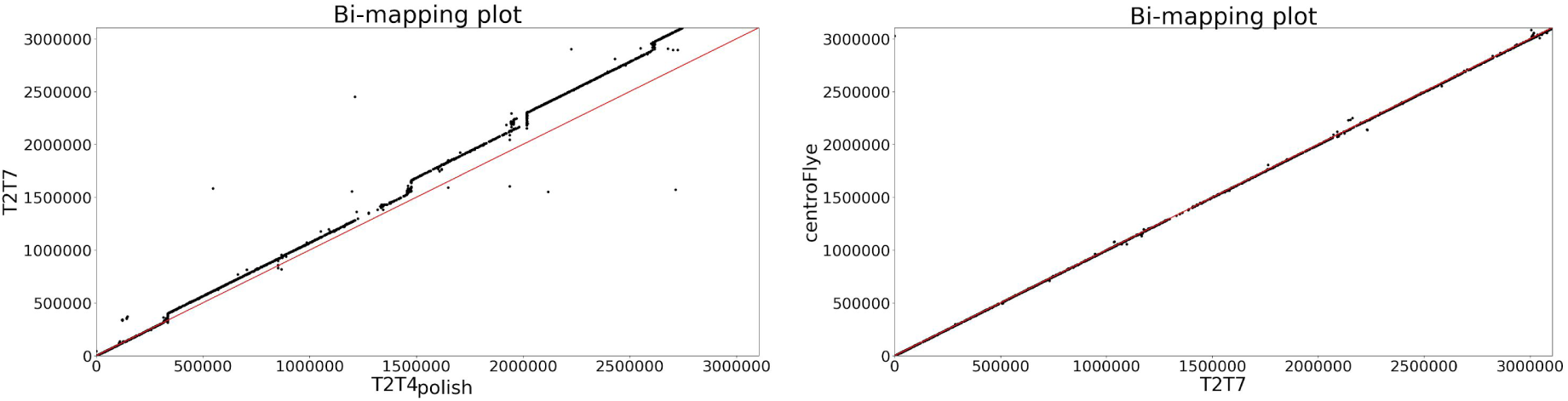
Bi-mapping plots for the T2T7 versus T2T4_polish_ and T2T7 versus centroFlye assemblies.

### Pairwise comparison of assemblies

Figure 13 shows bi-mapping plots for each pair of assemblies. As expected from the analysis of the breakpoint metric (Figure 9), the centroFlye and T2T7 assemblies are nearly identical. The T2T4_polish_ assembly differs from the T2T7 assembly around ∼350 kb, ∼1600 kb, ∼2100 kb, and ∼2800 kb (coordinates are given for the T2T7 assembly).

## Discussion

We presented the tandemMapper and tandemQUAST tools and applied them to various cenX assemblies. Although these tools detect flaws in ETR assemblies and provide a possibility to assess their quality, they have certain limitations discussed below.

### False assembly errors

TandemQUAST is based on mapping reads to the assembly and subsequent analysis. Such an approach implies that inherent errors or systematic biases in the sequencing platforms may affect evaluation of the assembly and bring in some discrepancies that could be considered as false assembly errors. To reduce this effect, tandemQUAST has an option of using accurate PacBio HiFi reads.

### Analysis of arbitrary ETRs in human and other genomes

Sequence and structural organization of ETRs, and particularly centromeres, varies widely across species. Since assembly of arbitrary ETRs remains an open problem, there is currently only one tool (centroFlye) for an automatic assembly of some ETRs and few examples of ETR assemblies. Thus, we purposefully limited the scope of our study to the recently completed human cenX assemblies. Since the Telomere-to-Telomere consortium aims to a gap-free assembly of a human genome that includes centromeric regions (Miga et al., 2019), we anticipate that more high-quality ETR assemblies will soon be generated. These new assemblies will help us in improving the tandemMapper and tandemQUAST tools.

### Analysis of diploid assemblies

Since centroFlye is now limited to haploid assemblies, the current version of tandemQUAST also focuses on haploid assemblies. Extending tandemQUAST functionality to diploid assemblies presents a complex algorithmic challenge. However, even effectively haploid cell lines may contain somatic heterogeneity due to clonal genomic instability in the cell culture. In this case, tandemQUAST can report heterozygous sites based on the discrepancies in mapped reads.

### Analysis of transposable elements in ETRs

TandemMapper currently masks TEs before selecting locally unique *k*-mers. This approach is not optimal for TE-rich centromeric regions such as Drosophila centromeres (Chang et al., 2019). We plan to minimize the influence of TEs on *k*-mer selection without masking them by setting a limit on the maximum number of *k*-mers that can be selected in each window of a fixed length (e.g., 5 kb).

### Using additional data types for assessing quality of ETR assemblies

We used accurate HiFi PacBio reads to analyze various centromere assemblies but not *bacterial artificial chromosomes* (*BACs*) and other alternative technologies that represent valuable resources for analyzing tandem repeats (see Appendix “Alternative technologies for ETR assembly quality assessment”).

For example, a BAC from an ETR is often easier to assemble than an entire long ETR such as a centromere. For example, centromere Y was recently sequenced using ONT reads to generate assemblies of BACs spanning this centromere (Jain et al., 2018a). However, certain limitations of the BAC technology make BACs a non-ideal option for ETRs sequence classification, (Miga et al., 2019). In particular, BACs (i) do not represent a high-throughput approach and thus limit the scope of studies, (ii) have severe differences in coverage that complicate analysis, (iii) require partial restriction digests that introduce biases in cloning, (iv) may have secondary structures making them incompatible with a bacterial host, (v) since existing short-read assemblers are unable to assemble highly repetitive centromeric BAC from short reads (or even Sanger reads), it is not clear how to reproduce the semi-manual assemblies of such BACs (some of them assembled two decades ago) with current state-of-the-art assemblers like SPAdes (Bankevich et al., 2012). It is also difficult to accurately assemble BACs from centromeres using long error-prone reads, e.g., recent large BAC sequencing effort has not resulted in assembling such BACs (Dennis et al., 2017). Thus, if a BAC sequence and a centromere assembly disagree, it is not clear whether this disagreement is caused by an error in the BAC assembly or an error in the centromere assembly. A possible way to address this challenge is a hybrid BAC assembly that combines short and long reads like in Jain et al., 2018a.

## Acknowledgements

We are grateful to Ivan Alexandrov for many insightful comments.

## Funding

This work was supported by St. Petersburg State University, St. Petersburg, Russia (grant ID PURE 28396291).

## Appendix: Unit-based statistic

If an assembly is represented as an array of monomers, tandemQUAST splits this array into repeated *units* (a sequence of monomers, e.g., a series of twelve monomers forming a HOR on cenX can be represented as *m*_*1*_*m*_*2*_…*m*_*12*_). To automatically derive a unit, tandemQUAST uses the StringDecomposer tool (Dvorkina et al., 2019) to translate the assembly from the nucleotide to the monomer alphabet (the alphabet size is the number of distinct monomers). Afterwards, it collects all *t*-mers in the monomer alphabet (the default value *t*=5), calculates the average distance *d* between two consecutive occurrences of the same *t*-mer, and selects the most frequent *d-*mer in the monomer alphabet as a *standard unit.* Afterwards, it removes all standard units and split the rest of the sequence into *non-standard units*, where each non-standard unit is the longest substring of a standard unit sequence. For example, given a standard unit *m*_*1*_*m*_*2*_*m*_*3*_*m*_*4*_*m*_*5*_*m*_*6*_*m*_*7*_*m*_*8*_*m*_*9*_*m*_*10*_*m*_*11*_*m*_*12*_, the monomer sequence *m*_*1*_…*m*_*9*_*m*_*5*_…*m*_*12*_ will be split into two units *m*_*1*_…*m*_*9*_ and *m*_*5*_…*m*_*12*_. TandemQUAST reports the assembly length in units, the number of distinct units, the number of monomers per each unit, and the unit frequency in the assembly and the read set.

Analysis of the *simulated*_*del_monomer*_ assembly demonstrated that it has 495 units, 494 of them are standard 12-monomers *m*_*1*_…*m*_*12*_ units, and, as expected, two units have non-standard sequences *m*_*1*_*m*_*2*_*m*_*3*_ and *m*_*7*_…*m*_*12*_.

Table S1 lists the distinct HOR units and their distribution in the assemblies and the reads. The centroFlye and T2T7 assemblies share the same set of units: 1536 HOR units, including 65 non-standard units. The centroFlye and centroFlye_polish_, as well as T2T4 and T2T4_polish_ assemblies also have the same set of units. The T2T4 assembly has a smaller length than the centroFlye and T2T7 (∼2.7Mbp vs ∼3.1Mbp), so the total number of units is lower, although the set of non-standard units is the same. All non-standard units are supported by reads.

**Table S1.**
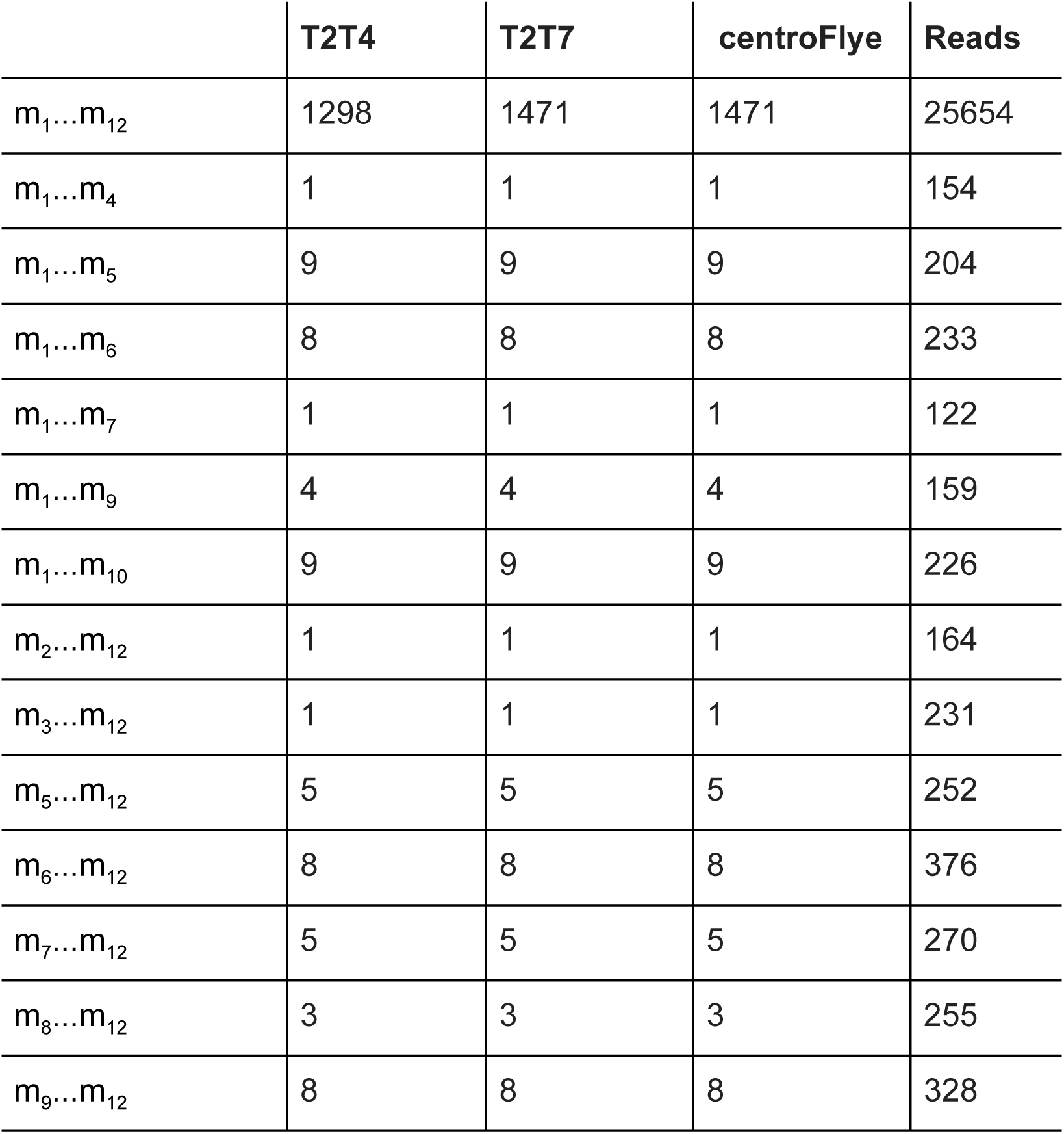
Distribution of distinct units in the T2T4, T2T7, and centroFlye assemblies and the read set. The first and the last units in the assembly are not listed in the table. The first unit in T2T4 and T2T7 assemblies is m_4_…m_12_, and in the centroFlye assembly is m_6_…m_12_. The last unit in all assemblies is m_1_…m_10_. The first unit in centroFlye assembly differ from those in T2T4 and T2T7 assemblies because of the choice of start sites and differences in the consensus HOR sequence.

|  | **T2T4** | **T2T7** | **centroFlye** | **Reads** |
| --- | --- | --- | --- | --- |
| $m_1...m_{12}$ | 1298 | 1471 | 1471 | 25654 |
| $m_1...m_4$ | 1 | 1 | 1 | 154 |
| $m_1...m_5$ | 9 | 9 | 9 | 204 |
| $m_1...m_6$ | 8 | 8 | 8 | 233 |
| $m_1...m_7$ | 1 | 1 | 1 | 122 |
| $m_1...m_9$ | 4 | 4 | 4 | 159 |
| $m_1...m_{10}$ | 9 | 9 | 9 | 226 |
| $m_2...m_{12}$ | 1 | 1 | 1 | 164 |
| $m_3...m_{12}$ | 1 | 1 | 1 | 231 |
| $m_5...m_{12}$ | 5 | 5 | 5 | 252 |
| $m_6...m_{12}$ | 8 | 8 | 8 | 376 |
| $m_7...m_{12}$ | 5 | 5 | 5 | 270 |
| $m_8...m_{12}$ | 3 | 3 | 3 | 255 |
| $m_9...m_{12}$ | 8 | 8 | 8 | 328 |

## Appendix: “Alternative technologies for ETR assembly quality assessment”

### CLR PacBio reads

probably add little to centromere assemblies since they are shorter than ONT reads and have similar error rates. Although they are better suited for polishing than ONT reads, difficulties with mapping shorter error-prone reads to repetitive centromeres may offset this advantage.

### Optical mapping

data was used by the T2T Consortium only for quality assessment (Miga et al., 2019). Even though incorporating optical mapping data into tandemQUAST remains an open problem, we hypothesize that the quality assessment metrics based on other data types, such as HiFi PacBio read, will be more beneficial.

### Hi-C data

Mapping of short Hi-C reads to ETRs presents a complex challenge that, to the best of our knowledge, remains unaddressed. Even though Hi-C data may be useful for quality assessment of ETR assemblies (especially for analysis of diploid assemblies) it is non-trivial to incorporate such data into tandemQUAST.

### 10X Genomics

data may potentially be useful but it is also non-trivial to incorporate this data type in tandemQUAST. We note that an even simpler problem of developing a 10X-based tool for analyzing quality of general assemblies remains unsolved.

